# A human-specific switch of alternatively spliced *AFMID* isoforms contributes to *TP53* mutations and tumor recurrence in hepatocellular carcinoma

**DOI:** 10.1101/169029

**Authors:** Kuan-Ting Lin, Wai-Kit Ma, Juergen Scharner, Yun-Ru Liu, Adrian R. Krainer

## Abstract

Pre-mRNA splicing can contribute to the switch of cell identity that occurs in carcinogenesis. Here we analyze a large collection of RNA-Seq datasets and report that splicing changes in hepatocyte-specific enzymes, such as *AFMID* and *KHK*, are associated with HCC patients’ survival and relapse. The switch of *AFMID* isoforms is an early event in HCC development, and is associated with driver mutations in *TP53* and *ARID1A*. Finally, we show that the switch of *AFMID* isoforms is human-specific and not detectable in other species, including primates. The integrative analysis uncovers a mechanistic link between splicing switches, *de novo* NAD^+^ biosynthesis, driver mutations, and HCC recurrence.

## Introduction

Liver cancer is the second leading cause of cancer death worldwide, and has very poor prognosis, with an incidence rate almost equal to the mortality rate (ratio = 0.95) (Ferlay et al. 2015). The global incidence of liver cancer has increased in the past 20 years, resulting in a doubling in disease-specific mortality (Llovet et al. 2015). Hepatocellular carcinoma (HCC) is the primary malignancy of the liver. The only approved drug for HCC is the protein tyrosine kinase inhibitor sorafenib, which can only prolong survival by about 3 months. Surgical resection has the best prognosis for long-time survival, but only a minority (~15%) of HCC patients have enough normal liver remaining at the time of diagnosis. Even if surgical resection is successful, most HCC patients (~90%) die within five years, because of intrahepatic recurrent HCC tumors (HCCs). The five-year survival rate is ~17% in the United States. Unfortunately, recent clinical trials of experimental HCC drugs have all failed (Llovet et al. 2015). Accordingly, there is an urgent unmet clinical need in prevention, diagnosis, prognosis, and treatment for this deadly cancer.

HCC cells are highly heterogeneous: different areas within the same tumor often have different patterns of morphology, immunohistochemical staining, and driver mutations (Friemel et al. 2015). The negative results from recent HCC clinical trials also highlight the intrinsic resistance of HCC to therapies (Villanueva and Llovet 2014). One important aspect of how cell identity is determined is through alternative pre-mRNA splicing patterns, which are regulated in a cell-type-specific manner. Recent studies identified some recurrent splicing events in HCC, but they did not establish associations with clinical information, and were limited to small patient cohorts (Sebestyen et al. 2015; Zhang et al. 2015; Sebestyen et al. 2016). Also, the detection methods used in the previous studies were limited to analyzing only two isoforms at a time.

To provide an integrative analysis of splicing patterns during the transition from hepatocytes to HCC cells, the present study analyzed ~6,000 samples of RNA-Seq data comprising human hepatocytes, Kupffer cells, adult and fetal livers, dysplastic lesions, early HCCs, HCCs, and cancer cell lines from various tissue types. We sought to identify robust splicing events associated with survival, recurrence, and driver mutations in HCC, by using a modified Percent Spliced-In (PSI) index (see Methods). In particular, we describe an *AFMID* alternative splicing event and propose that it plays a critical role in early HCC development and progression.

## Results

### Concordant splicing events in HCC tumors and liver-cancer cell lines

Liver-cancer cell lines are malignant clones derived from heterogeneous liver tumors. Concordant
splicing events that coexist in both HCCs and liver-cancer cell lines could be among the main characteristics preserved in liver-cancer evolution. Concordance means that the splicing event involves the same exon/exons, and increases or decreases in the same direction. To identify splicing events, we used a new PSI index to analyze RNA-Seq datasets (Methods and Supplementary Fig. S1). We started with an RNA-Seq dataset of 11 primary HCCs and matched adjacent normal liver (ANL) tissues from a recent study (Jhunjhunwala et al. 2014). Also, we analyzed 136 non-HCC liver samples from the Genotype-Tissue Expression (GTEx) database, and 16 liver-cancer cell lines (Consortium 2015; Klijn et al. 2015). We identified 436 and 1,992 robust splicing events in HCCs and liver-cancer cell lines, respectively (Supplementary Table S1 and S2). To identify highly reproducible splicing events, we required at least 20 supporting reads for the event in at least 80% of the samples.

Among the splicing events, we identified 136 overlapping events involving 135 alternative exons with concordant splicing changes (Fig. 1A). The top 10 genes with the largest PSI changes were *PEMT*, *KIAA1551*, *NUMB*, *FN1*, *MYO1B*, *USO1*, *RPS24*, *AFMID*, *KHK*, and *ARHGEF10L* (Fig. 1B). *PEMT* (phosphatidylethanolamine N-methyltransferase) mainly expresses the *PEMT_B_* isoform (NM_007169) in ANLs, but we found that it switches to greater expression of the *PEMT_A1_* isoform (NM_148172) in HCCs (Supplementary Fig. S2) (Shields et al. 2001). *NUMB* is a cell-fate determinant in cell development; the *NUMB_L_* isoform is known to be up-regulated in HCCs (Lu et al. 2015). *FN1* (fibronectin 1) had increased inclusion of its EDB exon and the function of the *FN1_EDB_* isoform was recently reported (Bordeleau et al. 2015). *USO1* (vesicle transport factor p115) showed a reduction in exon 13 inclusion (Chr4:76716488-76716509) and was recently proposed as a potential splicing marker in HCC (Danan-Gotthold et al. 2015). *KHK* (ketohexokinase) switched from *KHK_C_* to *KHK_A_* isoforms. A recent study showed that switching *KHK_A_* to *KHK_C_* can induce heart disease (Mirtschink et al. 2015). Conversely, switching *KHK_C_* to *KHK_A_* drives HCC development (Li et al. 2016). *AFMID* (arylformamidase) showed a decrease in the full-length isoform, and a higher proportion of the other two alternative isoforms in liver-cancer cells; one isoform skips five exons (exon 5 to exon 9) and the other skips four exons (exons 5, 7, 8, and 9). Among these top 10 genes, PEMT, KHK and AFMID are liver-specific enzymes. The events involving the alternative exons of *AFMID* had the most significant p-values (Fig. 1C). Exon 6 of *AFMID* has both increased and decreased PSI values in liver-cancer cells because it is present in the two full-length isoforms (*AFMID_FL1_* and *AFMID_FL2_*), and in an alternative isoform (*AFMID_e6_*), that skips exons 5, 7, 8, and 9 (Fig. 1D). In liver-cancer cells, the exon 6 PSI values were increased for the *AFMID_e6_* isoform, and decreased for the *AFMID_FL1_* and *AFMID_FL2_* isoforms.

**Figure 1.**
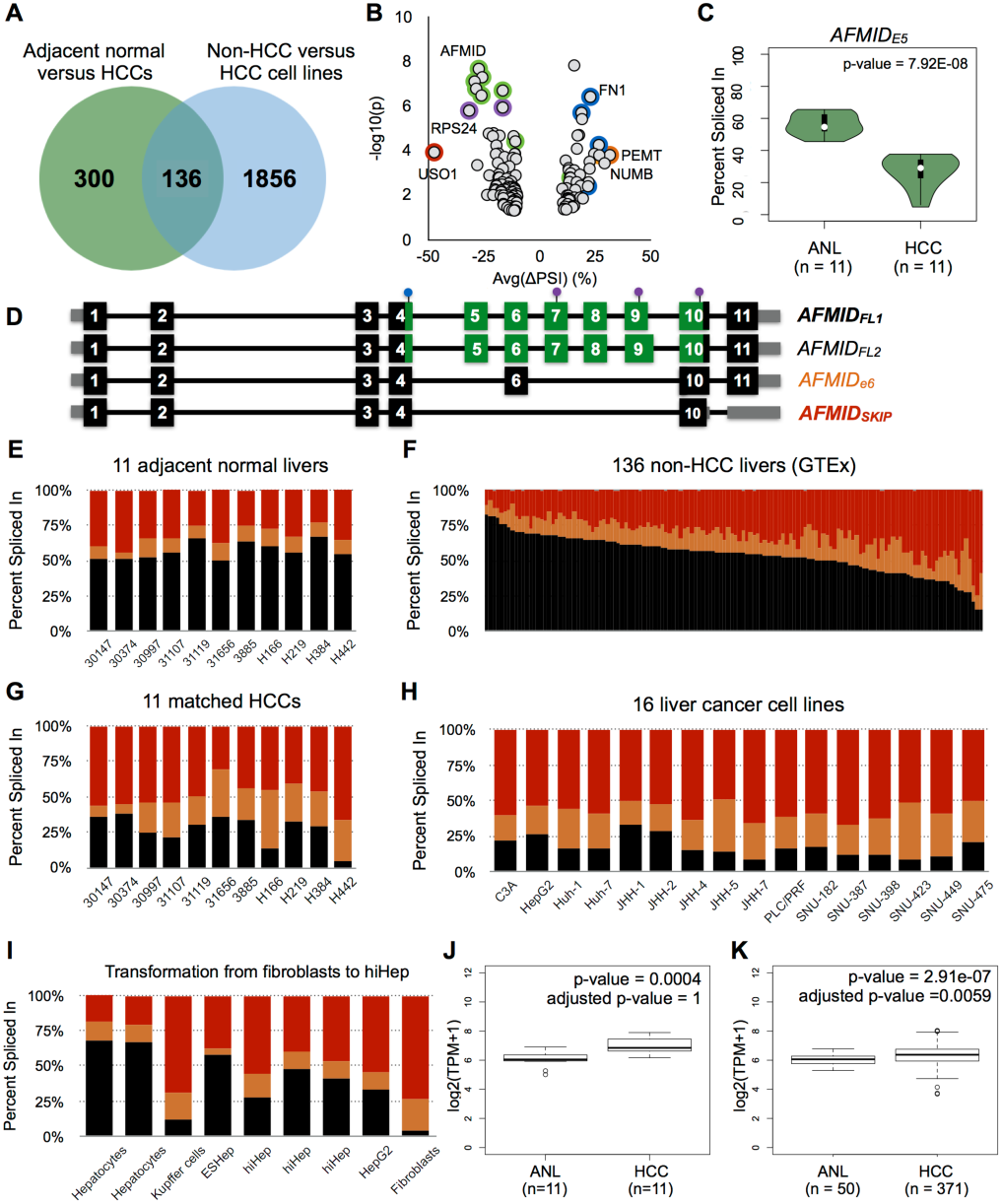
Concordant splicing events in HCC patient samples and liver-cancer cell lines. **(A)** The Venn diagram shows the number of overlapping splicing events between two independent comparisons: 11 adjacent normal livers (ANLs) versus 11 matched HCCs (green circle); and 136 non-HCC liver samples versus 16 liver-cancer cell lines (blue circle). **(B)** Each dot in the dot plot represents an alternative exon. The x-axis shows the average PSI change of an alternative exon between ANL and HCCs. The y-axis is the statistical significance, denoted as –log_10_(p), of the splicing event in 11 ANLs versus 11 HCCs. **(C)** The vioplot shows the PSI distributions of exon 5 of *AFMID* in 11 ANLs and 11 matched HCCs. **(D)** Exon/intron structure of *AFMID* isoforms. Each dark block is a coding exon; the gray boxes represent the UTR exons or portions of exons; the green boxes denote domain regions. Three purple dots represent the active-site residues (Ser164, Asp247, and His279). The blue dot on exon 4 represents the HGGXW motif (the oxyanion hole). **(E, F, G, and H)** The stacked bar charts in **(E)** and **(G)** represent the estimated proportion of *AFMID* isoforms in the 11 pairs of ANLs and matched HCCs, based on RNA-Seq junction reads. **(F)** shows the 136 non-HCC liver samples, and **(H)** shows the 16 liver-cancer cell lines. **(I)** The stacked bar chart shows the PSIs of *AFMID* isoforms in human hepatocytes, Kupffer cells, hiHep cells (ESHep and hiHep), HepG2, and skin fibroblasts. **(J and K)** The two box plots show the overall gene-expression level of *AFMID* in 11 matched HCC patient samples and 371 unmatched HCC patient samples from TCGA. Expression levels are shown as transcripts per million (TPM).

### The switch of *AFMID* isoforms corresponds to loss of normal hepatocyte identity

*AFMID* is located on Chromosome 17q25.3 and encodes arylformamidase, a controlling enzyme in tryptophan metabolism. *AFMID* expression level is evolutionarily constrained across multiple species (Pervouchine et al. 2015). *AFMID* generally expresses four isoforms, including *AFMID_FL1_*, *AFMID_FL2_*, *AFMID_e6_*, and *AFMID_SKIP_* (Fig. 1D). *AFMID_FL1_* is the major isoform and has a shorter exon 9 than *AFMID_FL2_*. In the present study, we used the PSI of exon 5 to represent the PSI of *AFMID_FL_*, as both *AFMID_FL1_* and *AFMID_FL2_* share the same exon 5. *AFMID_FL_* has a HGGXW motif (in exon 4), an alpha/beta hydrolase-fold domain (in exons 4 to 10), and an active site triad (in exons 7, 9, and 10, respectively) (Fig. 1D) (Pabarcus and Casida 2002; Pabarcus and Casida 2005). In normal or non-HCC livers, *AFMID* primarily expresses *AFMID_FL_* (Fig. 1E and 1F). In HCCs or liver-cancer cell lines, *AFMID* expresses mostly *AFMID_SKIP_* and *AFMID_e6_* (Fig. 1G and 1H). *AFMID_SKIP_* splices out exons 5 to 9, where the core-domain region resides. We also found that human hepatocytes had the highest PSI values of *AFMID_FL_*, and hiHep cells had higher PSI values of *AFMID_FL_* than HepG2 and fibroblasts (Huang et al. 2014). The *AFMID_SKIP_* and *AFMID_e6_* isoforms are the dominant isoforms in human Kupffer cells (Fig. 1I) (Costa-Silva et al. 2015). Both sets of data showed that the high-*AFMID_FL_* pattern is characteristic of human hepatocytes. Interestingly, although the PSI values of *AFMID_FL_* were significantly decreased in HCCs, the overall gene-expression level of *AFMID* was maintained at similar levels between ANLs and HCCs (Fig. 1J and 1K). Real-time RT-PCR experiments with RNA from 20 ANLs and 19 HCCs showed that the overall *AFMID* level did not significantly change (p=0.2963), but the *AFMID_FL_* isoform level was significantly down-regulated in HCCs by about 2-fold (p=0.0042) (Fig. S5).

To determine the complete structure of *AFMID_SKIP_*, we analyzed PacBio long reads derived from several cancer cell lines (Tilgner et al. 2014). We found that most of the isoforms that lack exons 5 to 9 have exons 1 through 4 in GM12878, GM12891, GM12892, and K562 cell lines (Supplementary Fig. S3). 79 of the 135 alternative exons identified above were present again in the set of 250 splicing events. *AFMID_FL_* also had decreased PSI values in HCCs from the LIHC dataset (Fig. 2A). Survival analysis showed that 32 of the 135 alternative exons had significant log-rank p-values in both overall and recurrence-free datasets (Fig. 2B and Supplementary Table S4). The top 5 genes were *AFMID*, *C16ORF13*, *SLAIN2*, *STRA13*, and *KHK*. *AFMID* exons had outstanding power at predicting patient survival. The PSI values of exons 5 and 6 had the strongest prognostic values in the RNA-Seq data of TCGA (Fig. 2B). The overall survival was positively correlated with the PSI values of *AFMID_FL_* in HCC (p=0.0011) (Fig. 2E). Patients with lower PSI values of *AFMID_FL_* died sooner (hazard ratio = 1.7087, p=0.0035), and tended to have a recurrence earlier (hazard ratio = 1.8822, p=3.60e-05) (Fig. 2C and 2D). For predicting HCC recurrence, *AFMID* is similar to *MKI67* (encoding the proliferation marker Ki-67) which had a log-rank p-value of 3.80e-05. 63 of the 64 low-*AFMID_FL_* patients died within 5 years. The median survival for low-*AFMID_FL_* patients was ~11.77 months (30 days per month), whereas for high-*AFMID_FL_* patients it was ~19.95 months.

**Figure 2.**
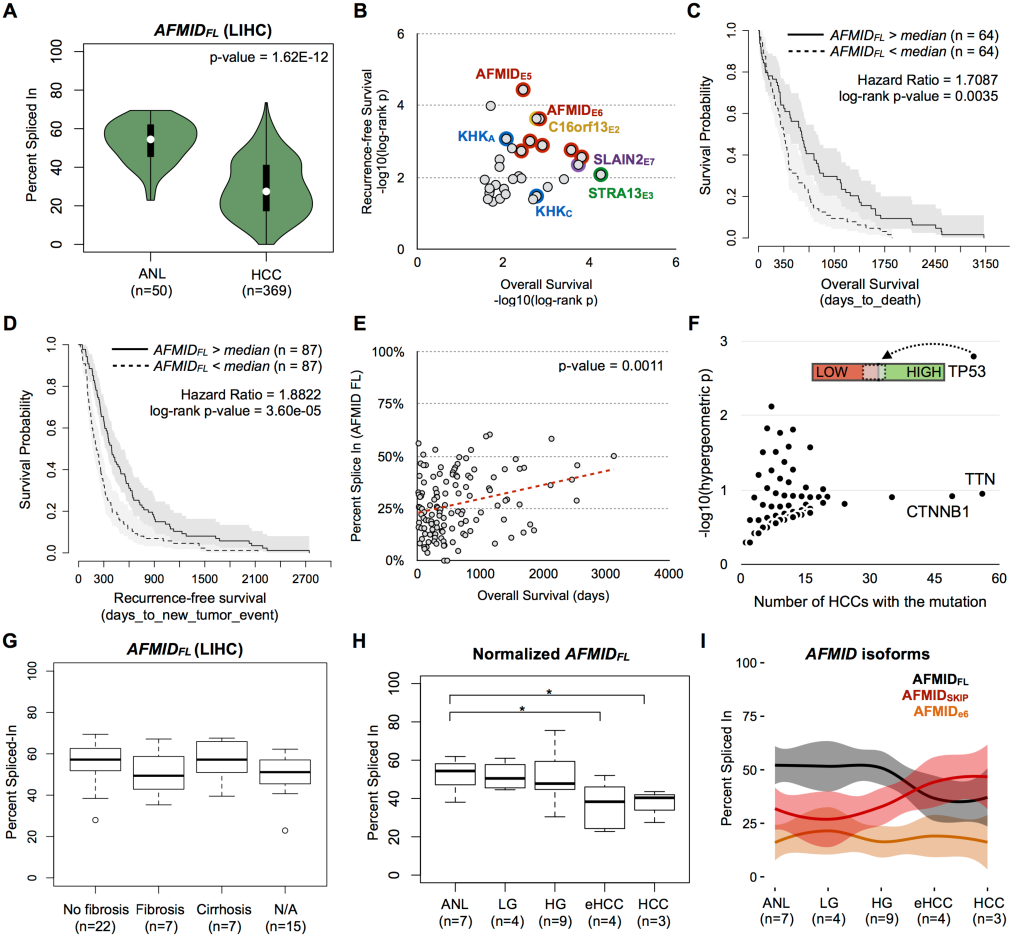
The switch of *AFMID* isoforms is associated with patient outcome and is already evident in early-stage HCC. (A) The vioplot shows the PSI distribution of *AFMID_FL_* isoform in adjacent normal livers (ANLs) and HCCs in the LIHC dataset of TCGA. **(B)** The dot plot summarizes the log-rank p values from overall (x-axis) and recurrence-free (y-axis) survival analysis, based on the LIHC dataset of TCGA. **(C and D)** The plots show the survival curves of *AFMID*-high and *AFMID*-low patients, based on the PSIs of the *AFMID_FL_* isoform in overall and recurrence-free survival analysis, respectively. The PSI of the *AFMID_FL_* isoform was determined by using the PSI of exon 5. **(E)** The dot plot shows the correlation between the PSI of the *AFMID_FL_* isoform and overall survival days. **(F)** The plot shows the enrichment (y-axis) of mutated genes in high-*AFMID_FL_* or low-*AFMID_FL_* HCCs, based on the LIHC dataset of TCGA. The x-axis shows the number of HCCs carrying the mutated gene. Each dot represents one gene mutated in at least one of the HCC samples. The bar labeled *TP53* demonstrates that there are more *TP53*-mutated HCCs (light-gray bar) in low-*AFMID_FL_* HCCs (red bar). **(G)** The box plots show the PSIs of the *AFMID_FL_* isoform in ANLs from patients with no fibrosis, fibrosis, or cirrhosis, in the LIHC dataset. N/A means that the annotation of the liver sample was not available. **(H)** The box plot shows the PSI distribution of the *AFMID_FL_* isoform in ANLs, low-/high-grade dysplastic lesions (LG/HG), early HCCs (eHCC), and HCCs. **(I)** The plot shows the PSI patterns of *AFMID_FL_* (black), *AFMID_SKIP_* (red), and *AFMID_e6_* (orange) isoforms in different groups of liver samples.

### The decrease of *AFMID_FL_* isoform is associated with driver mutations

The major source of nicotinamide adenine dinucleotide (NAD^+^) production in the hepatocyte is through tryptophan metabolism. A recent study showed that inhibition of the *de novo* NAD^+^ biosynthesis pathway leads to NAD^+^ depletion, DNA-damage responses, and HCC development in mice (Tummala et al. 2014). Feeding the mutant mice with nicotinamide riboside (NR), the precursor of the salvage pathway for generating NAD^+^, compensates for the loss of *de novo* NAD^+^ biosynthesis and prevents HCC development (Tummala et al. 2014). The study also showed that AFMID protein is down-regulated or not detected in human HCCs by western blotting, and depletion of *Afmid* in non-tumorigenic mouse liver cells (AML-12) resulted in aggressive tumors (Tummala et al. 2014). If the switch of *AFMID* isoforms increases DNA-damage responses in normal hepatocytes, low-*AFMID_FL_* HCCs would have a higher chance of accumulating driver mutations, such as in *TP53* and *ARID1A* (Villanueva and Llovet 2014). To test this hypothesis, we used the non-silent mutations. Among 369 HCC samples from TCGA, we found that 37 out the 54 TP53-mutated HCC samples were enriched in low-*AFMID_FL_* HCC samples (hypergeometric p=0.0016, Fig. 2F, Supplementary Table S5). In other words, low-*AFMID_FL_* HCC samples appear to have a 2-fold higher chance of gaining *TP53* mutations. Among 9,762 genes mutated in at least one of the 369 HCC samples, only *TP53* had a p-value lower than 0.01. *TTN* and *CTNNB1* were mutated in a similar number of HCC samples, but the p-values were not significant. Incorporating silent mutations yielded the same enrichment for *TP53*. 40 of 61 HCC samples with *TP53* mutations were enriched in the low-*AFMID_FL_* group (p=0.0016, Supplementary Table S6). In addition, we tested whether the 1st quartile (Q1) and the 4th quartile (Q4) of HCCs are associated with non-silent mutations in terms of PSI values of *AFMID_FL_*. Ranked by the PSI values of *AFMID_FL_*, 93 HCCs were in Q1 (PSI > 41%, high *AFMID_FL_*) and the other 93 HCCs were in Q4 (PSI < 17%, low *AFMID_FL_*). 26 of the 186 HCCs had *TP53* mutations, and 20 of them were enriched in Q4 (p=0.0020). Among the 6,370 mutated genes in the 186 HCCs, *ARID1A* also showed significant enrichment in Q4 (p=0.0289, Supplementary Table S7). Seven out of 8 HCCs with *ARID1A* non-silent mutations were enriched in Q4. Overall, there is a consistent enrichment of *TP53* mutations in low-*AFMID_FL_* HCCs.

### The switch of *AFMID* isoforms occurs in early-stage HCC

To establish when the switch of *AFMID* isoforms is likely to occur, we investigated the PSI distributions of *AFMID_FL_* in two datasets: (1) the LIHC datasets from TCGA; and (2) an RNA-Seq dataset that covers several stages in HCC development. First, among the 50 ANL samples from TCGA, 22 of the HCC patients showed no fibrosis, 7 showed fibrosis, and 6 showed cirrhosis. The PSI distributions of *AFMID_FL_* were not statistically different (Fig. 2G), indicating that the switch of *AFMID* isoforms is not associated with fibrosis or cirrhosis. Next, we investigated the other RNA-Seq dataset from a recent study (Marquardt et al. 2014). This dataset is composed of 7 ANL samples, 4 low-grade dysplastic lesions, 9 high-grade dysplastic lesions, 5 early HCCs, and 3 late HCCs. The sequencing depth for the samples in the RNA-Seq dataset ranges from 7 million to 339 million reads. This RNA-Seq dataset has particularly strong enrichment for reads in the 3’ end of *AFMID*. For example, in the 419 TCGA patient samples (50 ANLs and 369 HCCs), the PSI values of exon 5 and exon 9 had high correlation (correlation = 0.85) (Supplementary Fig. S6A). However, in the 28 patient samples, the PSI values of the two exons were weakly correlated (correlation = −0.16) (Supplementary Fig. S6B). The PSI values of exon 5 are generally much lower than the PSI values of exon 9. The imbalance in PSI values leads to inaccurate estimations of the *AFMID* isoform proportions if we use the PSI values of exon 5 alone to represent *AFMID_FL_*. Accordingly, we normalized the PSI of *AFMID_FL_* by using the average PSI of exon 5 and exon 9 in the second dataset. After normalization, the PSI distributions of *AFMID_FL_* were not statistically different in ANL, low-grade and high-grade dysplastic lesions. In contrast, the PSI values of *AFMID_FL_* were significantly lower in early HCCs (p= 0.0318) and HCCs (p=0.0356) than in ANLs (Fig. 2H). Combining the three isoforms in one figure, we could show that the PSI values of *AFMID_FL_* and *AFMID_SKIP_* start to intersect at the early stage of HCC (Fig. 2I).

### The decrease of *AFMID_FL_* isoform in other cancers

*AFMID* has highest expression in the liver, because of this organ’s high demand for NAD^+^ (400-800 μmol/kg protein) (Houtkooper et al. 2010). To investigate if cancers from other organs have the same switch, we analyzed RNA-Seq data from 5,213 samples from the GTEx portal, and from 675 cancer cell lines originated from 31 tissue types (Consortium 2015; Klijn et al. 2015). We found that all of the cancer cell lines expressed higher proportions of *AFMID_SKIP_* and *AFMID_e6_* isoforms (Fig. 3A). The 5,213 non-cancer samples generally had higher PSI values for the *AFMID_FL_* isoform, compared to their cancer cell line counterpart (Fig. 3B). Among the 15 matched tissue types, liver and kidney had the largest decrease in PSI values, on average, and lung had the most significant p-value (Fig. 3C). On the other hand, brain had very little change in PSI values. We note that the non-HCC livers with a lower RNA integrity (RIN) score in the GTEx dataset tend to have lower PSI values of *AFMID_FL_* (p=9.0e-06).

**Figure 3.**
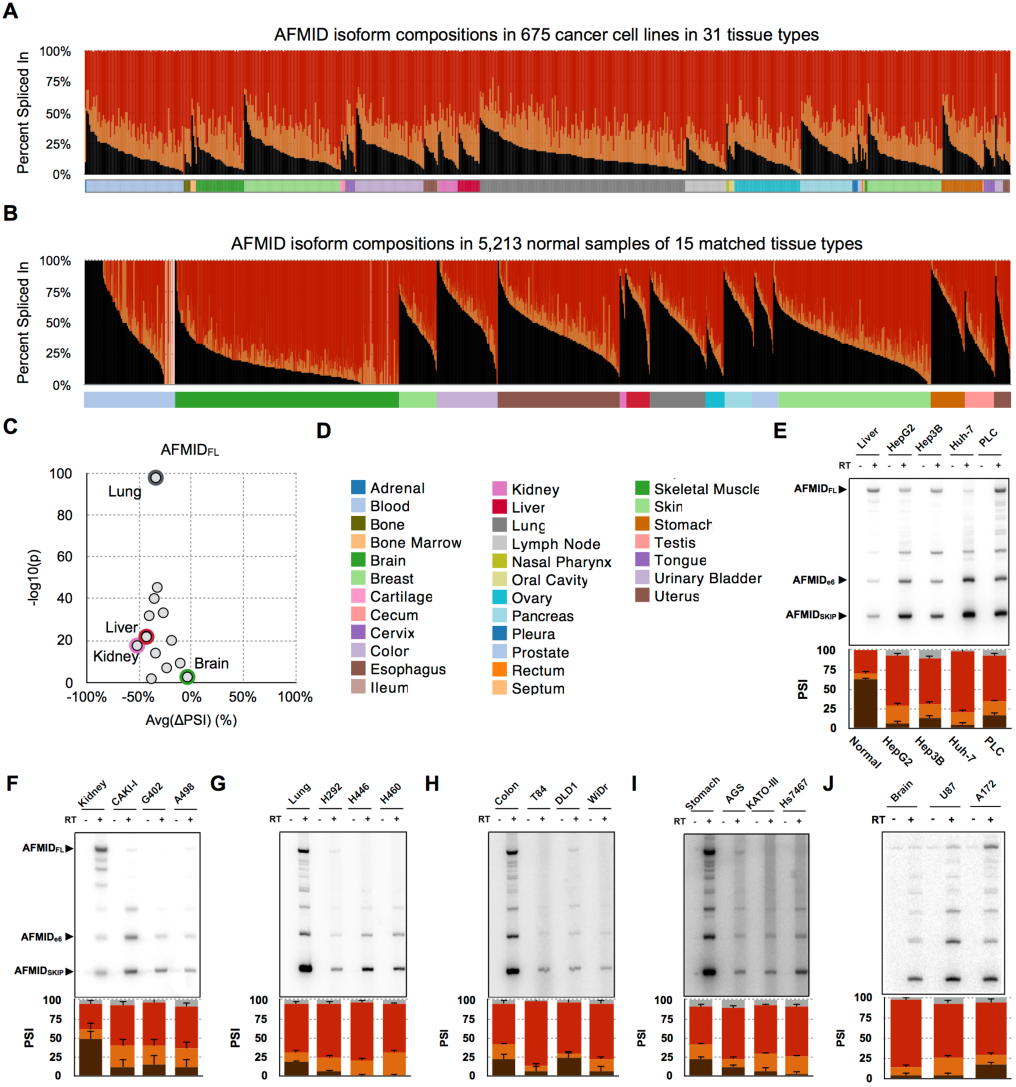
The switch of *AFMID* isoforms in cancers. **(A)** The stacked bar chart shows the proportions of *AFMID* isoforms (*AFMID_FL_*: black, *AFMID_SKIP_*: red, and *AFMID_e6_*: orange) in 675 cancer cell lines, based on RNA-Seq analysis. **(B)** The stacked bar chart shows the proportion of *AFMID* isoforms in non-cancer samples of 15 matched tissue types. **(C)** The dot plot shows the average ΔPSI between non-cancer samples and cancer cell lines from the same tissue type. The x-axis is the average ΔPSI, and the y-axis shows the statistical significance (-log_10_(p-value)) of the ΔPSI. **(D)** The color key of tissue types of **(A)** and **(B)**. **(E to J)** The six panels show radioactive RT-PCR results of *AFMID* isoforms in normal tissues and cancer cell lines from liver, kidney, lung, colon, stomach, and brain, respectively (24 amplification cycles). The three arrowheads on the left indicate the expected sizes of *AFMID_FL_*, *AFMID_e6_*, and *AFMID_SKIP_* isoforms, respectively. The stacked bar chart below each RT-PCR plot shows the average PSI values and standard deviations from triplicate experiments. The PSI bars of *AFMID_FL_*, *AFMID_e6_*, *AFMID_SKIP_*, and unknown bands are colored in black, orange, red, and gray, respectively.

To experimentally validate the event in multiple cancer types, we performed radioactive RT-PCR of RNA from normal liver, kidney, lung, colon, stomach, and brain tissue samples, versus their cancer cell line counterparts (Fig. 3E to 3J). We confirmed that *AFMID_FL_* is the dominant isoform in normal liver and kidney. We also confirmed that *AFMID_FL_* decreases and *AFMID_SKIP_* increases in most of the cancer cell lines tested, except for the brain cell lines (Fig. 3E to 3J). Real-time RT-PCR gave similar results. The overall *AFMID* expression levels were not significantly different between normal liver tissues and HepG2 and Hep3B cells, whereas Huh-7 and PLC/PRF/5 cell lines had significantly lower overall *AFMID* levels (Supplementary Fig. S7A). *AFMID_FL_* was generally lower in the four liver-cancer cell lines, whereas *AFMID_e6_* and *AFMID_SKIP_* were mostly unchanged (Supplementary Fig. S7A). Further RT-PCR analysis showed that human fetal liver also switched to the *AFMID_SKIP_* isoform (Supplementary Fig. S7B). The pattern is consistent with recent RNA-Seq data from human fetal liver (Gerrard et al. 2016).

### The switch of *AFMID* isoforms is human-specific

*AFMID* and *TK1* are one of the rarest anti-regulated head-to-head pairs conserved in many species (Li et al. 2006). *TK1* is thymidine kinase 1, whose expression levels fluctuate depending on the cell-cycle stage. Using an ultra-deep RNA-Seq dataset, we found zero supporting junction reads for *AFMID_SKIP_* and *AFMID_e6_* isoforms in fetal (E18), post-natal day 14 or day 28 (PN14 and PN28) and 3-month-old adult (A3M) mouse liver samples (Bhate et al. 2015). Likewise, we did not find supporting junction reads for *AFMID_SKIP_* and *AFMID_e6_* isoforms in RNA-Seq data from the *Mst1*^-/-^; *Mst2*^Flox/Flox^ mouse HCC model (Fitamant et al. 2015). Using RNA-Seq data from chimpanzee (*Pan troglodytes* and *Pan paniscus*), *Pongo pygmaeus*, *Macaca mulatta*, gorilla, mouse, and chicken, we again found that neither *AFMID_SKIP_* nor *AFMID_e6_* isoforms are expressed in liver, kidney, heart, muscle, and brain (Brawand et al. 2011; Barbosa-Morais et al. 2012). We conclude that *AFMID* splicing regulation is human-specific (Fig. 4A). Further radioactive RT-PCR showed no bands for the alternative isoforms in mouse liver and tumor samples (Fig. 4B). This analysis again confirmed that the alternative splicing regulation of *AFMID* isoforms is specific to human cells. From the mouse E18 versus PN28 comparison, we identified 2,149 splicing events involving 1,958 alternative exons. The ΔPSIs of 130 alternative exons from our RNA-Seq analysis were highly correlated with the ΔPSIs estimated by RT-PCR in the original paper (correlation = 0.8741) (Supplementary Fig. S8) (Bhate et al. 2015).

**Figure 4.**
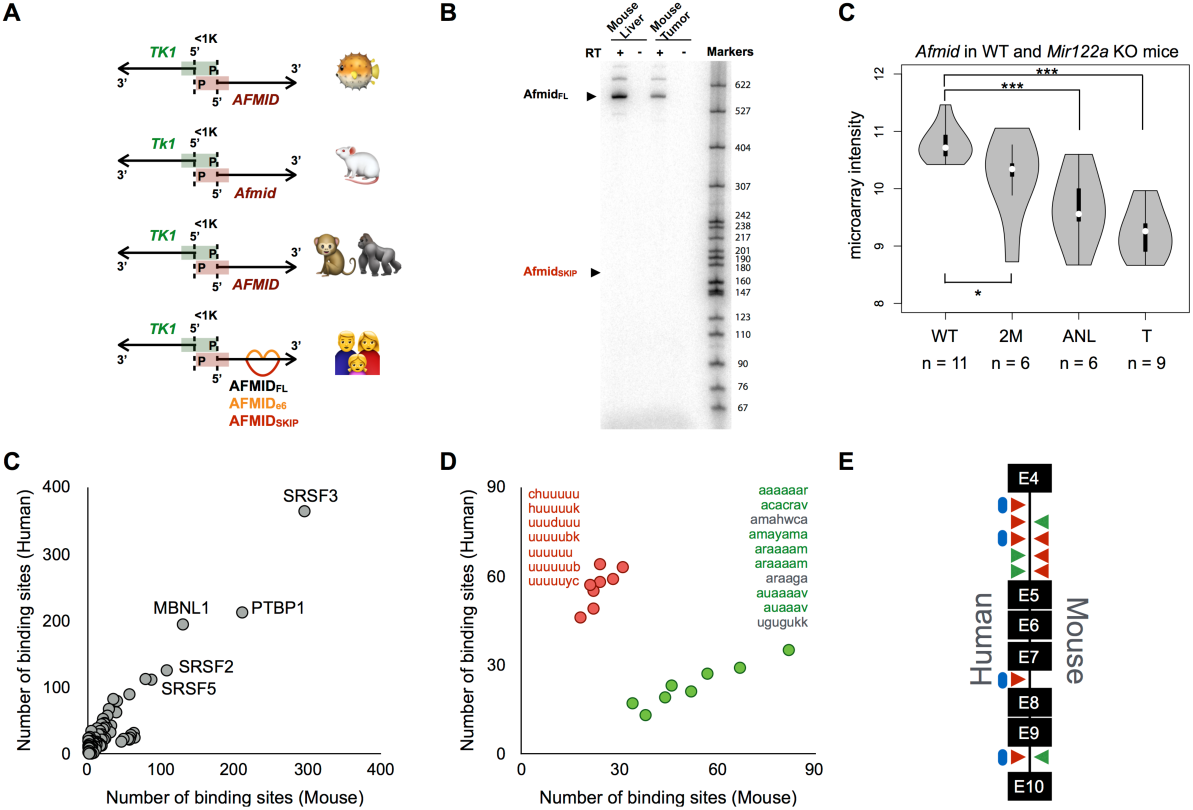
Phylogenetic comparisons of *AFMID* splicing patterns. **(A)** The bidirectional gene pair, *TK1* and *AFMID*, is a conserved structure in Fugu, mouse, monkey, gorilla, and human (from top to bottom). The black arrows indicate the direction of transcription for each gene. The promoter region is indicated by green and red bars for *TK1* and *AFMID*, respectively. The distance between the two transcription start sites is less than 1Kb. In the human diagram, the red arc indicates the *AFMID_SKIP_* isoform, and the orange arcs represent the *AFMID_e6_* isoform. (B) Radioactive RT-PCR analysis of *Afmid* isoforms in mouse liver and tumor samples. Expected sizes of isoforms are labeled on the left. Size markers are shown on the right. **(C)** The vioplot shows the expression patterns of *Afmid* in *Mir122a* knockout (KO) mice. WT: wild-type mice; 2M: 2-month-old *Mir122a*-KO mice that lack tumors; ANL: adjacent normal liver samples from older *Mir122a*-KO mice that have developed liver tumors; T: tumor part of the liver tumors from older *Mir122a*-KO mice. Two sample t-tests were used to determine the p-values. **(C)** The dot plot shows the number of presumptive binding sites of RNA-binding proteins in mouse (x-axis) and human (y-axis). Each dot corresponds to an RNA-binding protein. **(D)** Only RNA-binding proteins with ≥2-fold change in the number of binding sites are shown; Proteins with more predicted binding sites in human are shown in red, and those with more binding sites in mouse are shown in green. The motifs of RNA-binding proteins are colored in the same manner. The motifs shown in gray are from RNA-binding proteins with undetectable liver expression, based on GTEx data. **(E)** The diagram shows where the red and green motifs are located within the region from exon 4 to exon 10 of *AFMID*. Each triangle points to the location of a cluster of motifs in the intron. The locations of green and red motifs in the human gene are shown on the left, and those of the mouse gene are shown on the right. Blue dots represent the binding sites of HNRNPC in human cells, based on enhanced cross-linking immunoprecipitation data from ENCODE.

In contrast to human, other species regulate *AFMID* transcriptionally, such that *AFMID* is down-regulated in proliferative states. We found that *AFMID* is down-regulated in both fetal liver and liver tumors in mice (Fig. 4B) (Hsu et al. 2012; Tsai et al. 2012; Bhate et al. 2015). The 5’ and 3’ splice sites of exons 4, 5, and 10 had similar calculated strengths between human and mouse (data not shown). We investigated the potential binding sites of 94 RNA-binding proteins, which might function as splicing activators or repressors, in the region from exon 4 to exon 10 of *AFMID* (human) and *Afmid* (mouse) (Paz et al. 2014). We found that SRSF3, PTBP1, MBNL1, SRSF2, and SRSF5 had the most binding sites, on average, in human and mouse (Fig. 4C). The number of binding sites of the top 5 RNA-binding proteins was similar between human and mouse (Supplementary Fig. S9). On the other hand, among the 39 RNA-binding proteins with more than 20 binding sites, two groups of proteins had at least a 2-fold decrease or increase in the number of binding sites between human and mouse (Fig. 4D). The first group had more predicted binding sites in human, and includes CPEB2 (chuuuuu), CPEB4 (uuuuuu), HNRNPC (huuuuuk), HNRNPCL1 (huuuuuk), RALY (uuuuuub), TIA1 (uuuuubk), U2AF2 (uuuuuyc), and ZC3H14 (uuuduuu). The proteins in the first group shared similar motifs, with a string of Us. HNRNPCL1 is not expressed in the liver, based on GTEx data. The first group of proteins have predicted binding sites in intron 4 and intron 9 of human *AFMID*, but this pattern is largely lost in mouse *Afmid* (Fig. 4E and Supplementary Fig. S10A). Conversely, the first group of proteins gained an additional two clusters of predicted sites in intron 4 near the 3’ splice site of mouse *Afmid* (Fig. 4E and Supplementary Fig. S10A). The second group of proteins includes BRUNOL5 (ugugukk), HNRNPL (acacrav and amayama), IGF2BP3 (amahwca), KHDRBS1 (auaaaav), KHDRBS3 (auaaav), PABPC1 (araaaam), PABPC4 (aaaaaar), PABPN1 (araaga), and SART3 (araaaam). BRUNOL5, IGF2BP3, and PABPN1 are not expressed in the liver, based on GTEx data. Unlike the proteins in the first group, the proteins in the second group share A-rich motifs. They gained new binding sites in mouse in intron 4 and intron 9 (Fig. 4E and Supplementary Fig. S10B). Enhanced cross-linking immunoprecipitation data from ENCODE also showed that HNRNPC binds to most of the predicted regions in human cells (blue dots in Fig. 4E). This is consistent with our predictions.

## Discussion

HCC’s heterogeneity is a challenge for developing advances in prognosis and treatment (Friemel et al. 2015; Llovet et al. 2015). We tried to overcome this challenge by characterizing the splicing events in liver-cancer cells. We report that hepatocyte-specific splicing patterns have outstanding power in predicting HCC recurrence. Especially, the *AFMID* splicing event is associated with the presence of early driver mutations, such as mutated *TP53* and *ARID1A*. The switch of *AFMID* isoforms represents a new regulatory step in tryptophan/kynurenine metabolism, and revealed the disruption of *de novo* NAD^+^ biosynthesis in hepatocytes in the early stages of HCC development. Low-*AFMID_FL_* HCCs tend to have a higher chance of carrying *TP53* mutations, but not *CTNNB1* mutations. This is consistent with the current understanding that mutated *CTNNB1* is a later event (Friemel et al. 2015). Only the link between mutated *TP53* and the switch of *AFMID* isoforms was preserved during HCC evolution. Indeed, 7 of 16 liver-cancer cell lines we analyzed lacked *TP53* mutations (Klijn et al. 2015). Therefore, the switch of *AFMID* isoforms can occur without *TP53* mutations. Together with the evidence from fetal liver and *Mir122a^-/-^* mouse liver, it is readily apparent that the switch of *AFMID* isoforms is an early event in HCC development. The switch may play an important role in early HCC evolution, because it increases the chance of accumulating driver mutations in HCC-initiating cells (Fig. 5A).

**Figure 5.**
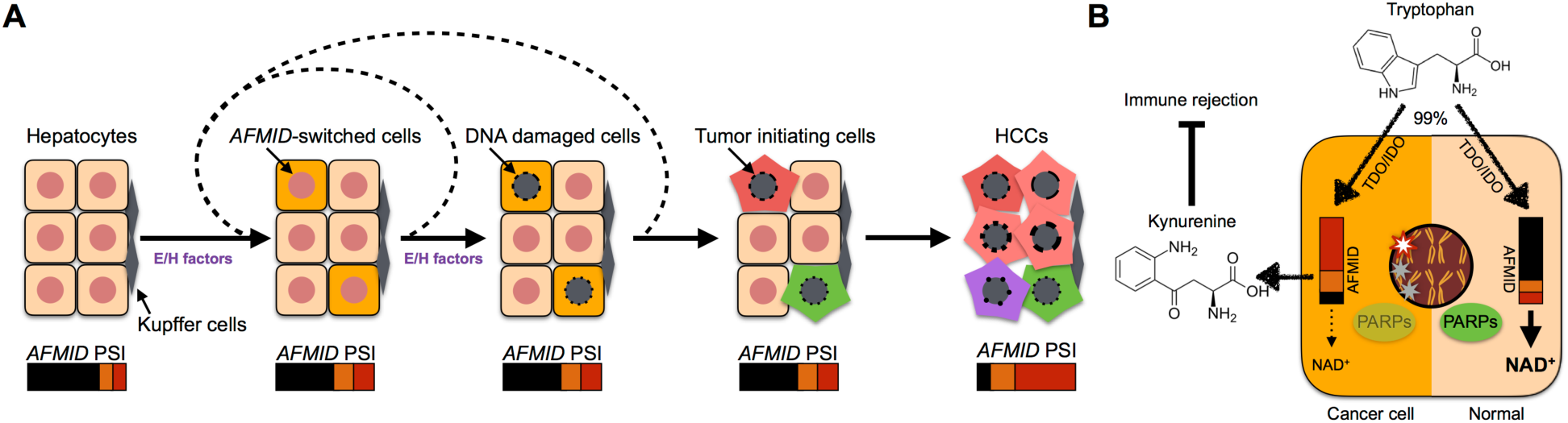
Model of the *AFMID* isoform switch in HCC. **(A)** The flow chart represents HCC progression (left to right). At the start, six representative hepatocytes are shown (nuclei in red). Later, because of environmental or hereditary factors (E/H factors), a subset of hepatocytes switches *AFMID* isoforms. The environmental factors include WNT signals in pericentral hepatocytes in daily liver regeneration, cytokines released during inflammation, chemical damage, and virus infection. The hereditary factors include driver mutations, such as in *TP53* and *ARID1A*. The E/H factors temporarily disrupt the identity of hepatocytes and reduce the NAD^+^ level in the hepatocytes. The reduced NAD^+^ level gives rise to increased DNA damage in the nucleus. After recursively accumulating driver mutations, the DNA-damaged hepatocytes become HCC initiating cells. The switch of *AFMID* isoforms can then be observed by bulk RNA-Seq. **(B)** The diagram shows two states of hepatocyte cells. On the right, a normal hepatocyte expresses high levels of *AFMID_FL_* isoform, and tryptophan can be converted to NAD^+^ for PARPs to fix DNA damage. On the left, the HCC cell expresses low levels of *AFMID_FL_* and has low NAD^+^, so DNA damage is increased and kynurenine is secreted to inhibit immune rejection.

NAD^+^ is a vital coenzyme in energy metabolism in eukaryotic cells (Houtkooper et al. 2010; Canto et al. 2015). NAD^+^ repletion increases life span in mice (Zhang et al. 2016). However, the NAD^+^/NADH ratio is very low in cancer cells; they maintain sufficient NAD^+^ for a high rate of glycolysis by converting pyruvate to lactate, while turning off other sources of NAD^+^ production (Liberti and Locasale 2016; Vander Heiden and DeBerardinis 2017). For example, the switch of *AFMID* isoforms impairs the major source of NAD^+^ production in hepatocytes. The switch may facilitate proliferation, but it also increases DNA-damage responses. For example, poly-(ADP-ribose) polymerase (PARP) and Sirtuin are both NAD^+^-dependent enzymes. PARP enzymes consume NAD^+^ to generate PAR polymers for repairing DNA. Sirtuin enzymes are associated with longevity, aging, and cancer (Herranz et al. 2010; Canto et al. 2015). Accordingly, the dysregulation of the *de novo* NAD^+^ pathway is a key event in HCC development. The switch of *AFMID* isoforms contributes to the accumulation of driver mutations and increases cancer susceptibility (Fig. 5B). Our discovery of the two human-specific isoforms (*AFMID_SKIP_* and *AFMID_e6_*) may lead to uncovering new mechanisms in tryptophan metabolism, as these are the predominant isoforms in cancer cells. Their roles in kynurenine secretion need to be further investigated. Switching the *AFMID_SKIP_* and *AFMID_e6_* isoforms back to *AFMID_FL_* may impact the secretion of kynurenine by redirecting the flux of tryptophan back to *de novo* NAD^+^ biosynthesis. This in turn may enhance NAD^+^ production, and reduce immune escape of cancer cells (Fig. 5B). Also, modulating the splicing switch has potential implications for neurodegenerative diseases (Vecsei et al. 2013).

In summary, the present study provides the first integrative analysis of splicing events in liver cancer. We identified new splicing-based biomarkers in hepatocyte-specific enzymes, such as PEMT, KHK, and AFMID. We found that *AFMID* alternative splicing constitutes a key event in liver carcinogenesis, and a new switch in tryptophan/kynurenine metabolism.

## Methods

### A new PSI index

Traditionally, the PSI index is denoted as *(a+b)/(a+b+2c)*, where ***a*** and ***b*** stand for the number of splice-junction reads connecting the alternative exon to the upstream and downstream constitutive exons, respectively (Barbosa-Morais et al. 2012). ***c*** stands for the number of junction reads connecting the two constitutive exons. The traditional equation is designed for simple splicing events with only one alternative exon, but it is ambiguous in the case of mutually exclusive exons, multi-exon skipping, and more complex events. Therefore, we modified the PSI index as follows:

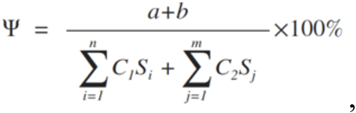

where ***C_1_*** and ***C_2_* stand for the upstream and downstream constitutive exons, respectively. *C_1_S_i_*** stands for the total number of junction reads whose 5’ splice site is connected to the upstream constitutive exon in a given splicing event. Similarly, ***C_2_S_j_*** stands for the junction reads whose 3’ splice site is connected to the downstream constitutive exon. Because the denominator is the sum of junction reads connecting to the flanking constitutive exons, the equation does not have the ambiguity for mutually exclusive exon events in which ***c*** might not exist. Also, in the new PSI equation, ***a*** and ***b*** stand for the number of junction reads connecting the alternative exon to its upstream and downstream exons, respectively; ***a*** and ***b*** do not necessarily reflect connections to ***C_1_*** and ***C_2_*** exons. For alternative splice site events, only ***C_1_*** or ***C_2_*** is used in the denominator, because the event only involves one constitutive exon.

The new PSI index is more flexible and can accurately compute PSI values of individual exons in more complex splicing events. For example, the splicing events involving mutually exclusive exons of *KHK* were not reported in previous HCC studies (Danan-Gotthold et al. 2015; Sebestyen et al. 2016). Also, single-exon PSI approaches can simply ignore multi-exon splicing events. For example, MISO cannot detect the multi-exon splicing events of *AFMID and MYO1B*(Katz et al. 2010). Moreover, previous methods failed to report the PSI values of *AFMID_e6_* (illustrated in Supplementary Fig. 1). Exon 6 of *AFMID* is used by both *AFMID_FL_* and *AFMID_e6_* isoforms, which can be detected by the new PSI index. Finally, the new PSI index can more accurately detect changes involving alternative 5’ splice sites or 3’ splice sites. For example, *PEMT* uses two alternative 5’ splice sites in *PEMT_A1_* and *PEMT_A2_*. Because the new PSI index takes into account all the junction reads involving the 3’ splice site of exon 4, the switch from *PEMT_B_* to *PEMT_A1_* can be accurately detected (Supplementart Fig. 2).

In the present study, the alternative exons were identified based on the Ensembl 75 gene annotation. For a given alternative exon, each sample was required to have more than 20 supporting junction reads in the denominator of the PSI index. If fewer than 80% of the samples met the criteria, the splicing events were not considered as candidates for highly reproducible splicing events. In addition, splicing events with lower than 10% PSI change or whose p-value was larger than 0.05 were also excluded.

### RNA-Seq data process

RNA-Seq datasets were downloaded from several sources, such as Sequence Read Archive (SRA), European Genome-phenome Archive (EGA), TCGA, and GTEx. The RNA-Seq dataset of the 11 primary HCCs and matched normal livers were downloaded from SRA and aligned by STAR (2.4.1c)(Dobin et al. 2013). The RNA-Seq datasets of 136 non-HCC liver samples and 675 cancer cell lines were downloaded from EGA and aligned by STAR. For TCGA’s LIHC dataset, we downloaded the alignment files from The Cancer Genomics Hub (http://cghub.ucsc.edu) and extracted the counts of junction reads from the alignment files. Recurrent HCCs (02A or 02B) in the LIHC dataset were excluded. The PSI values of *AFMID* isoforms in the LIHC datasets were based on TCGA’s alignment results, which were processed using MapSplice(Wang et al. 2010). The PSI values of *AFMID_FL_* from TCGA’s alignment files in 8 randomly selected HCCs were almost identical to the PSI values based on the alignment files by STAR (correlation = 0.9983). In addition, the counts of junction reads of the 5,213 non-cancer samples in Fig. 4B were downloaded from the GTEx portal (http://www.gtexportal.org/). In summary, the present study used three different approaches to obtain splicing changes (Supplementary Fig. 1). GRCh37 (hg19) and Ensembl 75 were the reference genome and gene annotation for human datasets, respectively. Mm10 was the reference genome for mouse datasets. PPYG2 was the reference genome for *Pongo pygmaeus*. CHIMP2.1.4 was the reference genome for *Pan troglodytes* and *Pan paniscus*. MMUL1.0 was the reference genome for *Macaca mulatta*. GorGor3.1 was the reference genome for *Gorilla gorilla*. Galgal4 was the reference genome for *Gallus gallus*. The genome indeces of STAR were built using the default options, and sjdbOverhang was set to 100.

### Statistical analysis

The two sample t-test was used to elaborate the significance of PSI differences between non-cancer and cancer cells. The log-rank test was used for survival analysis. The hypergeometric test was used for enrichment analysis of somatic mutations. Correlation testing was based on Pearson’s product moment correlation coefficient. The adjustment method for p-values used the Bonferroni correction.

### RT-PCR

Total RNA was extracted from cell lines using Trizol (Invitrogen). Genomic DNA was removed by treatment with DNase I (Promega). Reverse transcription of 0.5 – 1 μg of total RNA was carried out using ImPromp-II reverse transcriptase (Promega). Semi-quantitative PCR in the presence of [α-32P]-dCTP was performed with Amplitaq polymerase (Applied Biosystems). The human-specific primer set (Forward: 5’-GGCCACCAGGAAGAGCCTGC-3’, Reverse: 5’-CCTTCTGGGTCAGATTCTCAAC-3’) was used to amplify endogenous *AFMID* transcripts; these primers anneal to exons 3 and 10. After 24 amplification cycles, the products were resolved using a 5% native polyacrylamide gel, and the resolved bands were visualized on a Typhoon 9410 phosphorimager (GE Healthcare). The signal intensities were quantified using ImageJ software(Schneider et al. 2012). Primer sequences are listed in Supplementary Table 8.

### Real-time PCR

0.5 μg of total RNA was extracted and reverse-transcribed as for RT-PCR. Complementary DNA
(cDNA) was analyzed on a 7900HT Fast Real-Time PCR system (ThermoFisher Scientific). Fold changes were calculated using the ΔΔCq method and are reported as three biological replicates with three technical repeats each with ± S.E.M. Real-time PCR results for HCC patient samples were obtained using a Bio-Rad system. For empirical validations, 20 ANLs and 19 HCCs from the Taiwan Liver Cancer Network were selected based on gender and cirrhosis status. These samples were used in accordance with the IRB procedures of Taipei Medical University. Primer sequences are listed in Supplementary Table 8.

## Footnotes

**AUTHOR CONTRIBUTIONS:** K.L. designed the study and performed the bioinformatics analysis. W.M. and S.J. performed radioactive RT-PCR experiments. Y.L. performed quantitative RT-PCR experiments for human HCC patient samples. K.L. and A.R.K. wrote the manuscript with input from all co-authors.

## ACKNOWLEDGMENTS

K.L., W.-K.M, J.S., and A.R.K. acknowledge support from NCI grant CA13106. We thank M. Wigler for sharing cancer cell lines, and D. Tuveson and D. Fearon for valuable suggestions. We thank the Taiwan Liver Cancer Network (TLCN) for providing the liver tissue samples.

